# Transcriptome analysis reveals novel regulators of the Scots pine stilbene pathway

**DOI:** 10.1101/2023.01.05.522892

**Authors:** Tanja Paasela, Kean-Jin Lim, Mirko Pavicic, Anni Harju, Martti Venäläinen, Lars Paulin, Petri Auvinen, Katri Kärkkäinen, Teemu H. Teeri

## Abstract

Stilbenes are developmentally induced metabolites in Scots pine heartwood where they have important role in protecting wood from decaying fungi. The stilbene pathway is also stress inducible, and ultraviolet (UV)-C radiation was among the first discovered artificial stress activators of the pathway. Many conditions that activate the pathway are known, but the specific transcriptional regulators and the biosynthetic enzyme responsible for activating the stilbene precursor cinnamate in the pathway are unknown.

Here we describe the first large-scale transcriptomic analysis of pine needles in response to UV-C exposure and treatment with translational inhibitor cycloheximide, both activating the transcription of stilbene pathway genes, to uncover the candidates for the transcriptional regulation of the pathway and the cinnamate activating CoA ligase. We show that the regulation of pine stilbene pathway has shared features with grapevine stilbene pathway. In both species, the stilbene pathway was transcriptionally activated after UV-C treatment, protein phosphatase inhibitors and plant hormones ethylene and jasmonate.

The pine stilbene synthase promoter retains its inducibility when transformed in plants that normally do not synthesize stilbenes. Pine stilbene synthase promoter was able to activate the expression of a reporter gene in Arabidopsis in response to UV-C exposure and phosphatase inhibitors, but not as response to plant hormone treatment. This indicates that stilbene synthase gene regulation occurs both via ancient stress-response pathway(s) but also via species specific regulators. With transcriptomic approach we identified candidate enzymes for cinnamate acting CoA ligase and transcription factors regulating the pathway.

## Introduction

Stilbenes are important secondary metabolites for both the induced and constitutive defense systems in many plant species (Chong et al., 2009). The stilbenes of Scots pine (*Pinus sylvestris* L.) consist mainly of pinosylvin (3,5-dihydroxystilbene) and its monomethylether (3-methoxy-5-hydroxystilbene). Additionally, a minute amount of dimethylether (3,5-dimethoxystilbene) can be found in pine heartwood (Hovelstad et al., 2006; Venäläinen et al., 2004). In comparison to angiosperm and spruce stilbenes, pine stilbenes are structurally different deriving from cinnamoyl-CoA instead of *p*-coumaroyl-CoA.

The biosynthetic pathway leading to stilbene formation consists of four enzymes. Stilbene synthase (STS) (Schwekendiek et al., 1992) and pinosylvin *O*-methyltransferase (PMT) (Paasela et al., 2017) are characterized, but the pathway-specific cinnamate activating CoA ligase is still unknown. In Aaron’s beard (*Hypericum calycinum*) and petunia (*Petunia hybrida*) cinnamate is activated by cinnamate:CoA ligase (CNL) in the biosynthetic pathway to benzoic acid and its derivatives (Klempien et al. 2012, Gaid et al. 2012). These enzymes have low sequence similarity with 4CL enzymes. Cinnamate can also be activated by a specific 4CL variant. In *Rubus idaeus* such variant has been discovered (Kumar & Ellis 2003) and in Arabidopsis (*Arabidopsis thaliana*) 4CL1-3 can activate cinnamic acid with low efficiency (Costa et al., 2005; Ehlting et al., 1999). It is not known either if there are stilbene pathway-specific phenylalanine ammonia lyases (PAL) or is it shared with flavonoid and/or lignin pathways.

We have previously studied the induction of stilbene biosynthesis under the developmental context of heartwood formation (Lim et al., 2016) and in response to mechanical wounding (Lim et al., 2021). The developmentally regulated heartwood stilbenes are important determinants of heartwood decay resistance (Belt et al., 2021; Harju & Venäläinen 2006). The pathway is also stress-responsive, and it has been shown that the stilbene pathway in seedlings is strongly induced by UV-C treatment (Schoeppner and Kindl, 1979; Preisig-Müller et al., 1999). UV-C represents the shortest wavelength region of the UV-spectrum, which is divided into UV-A (315-400 nm), UV-B (280-315 nm) and UV-C (100-280 nm). The most damaging wavelengths, including UV-C, are completely absorbed by the atmosphere from solar UV radiation, but it can be used as an experimental tool to study, for instance, programmed cell death (PCD), DNA damage and repair mechanisms (Danon et al., 2004; Gallego et al., 2000; Gao et al., 2008; Gentile et al., 2003) and stilbene pathway activation and regulation (Vannozzi et al., 2012; Xi et al., 2014; Suzuki et al., 2015). When used experimentally, UV-C produces a strong reactive oxygen species (ROS) burst mainly from chloroplasts and mitochondria (Gao et al., 2008). UV-C responses are mediated by nonspecific stress signaling pathways originating from ROS, DNA damage and wound/defense signaling involving ethylene, jasmonic acid (JA) and salicylic acid (SA) (Jenkins, 2009; Nawkar et al., 2013). The effect of UV-C in activating the stilbene pathway has been studied more in detail in grapevine (*Vitis vinifera* L.) (Vannozzi et al., 2012; Xi et al., 2014; Suzuki et al., 2015).

It is known that stilbene biosynthesis in pine is transcriptionally regulated, but the mechanisms and components of the signaling pathways under different inductive conditions are unknown. Grapevine produces the stilbene resveratrol in response to abiotic and biotic stress, and during fruit development. It has been shown that the plant hormones JA, its volatile derivative methyl jasmonate (MeJA) and ethylene activate the stilbene synthase encoding genes (*VvSTS*) in grapevine (Belhadj et al, 2008; Grimmig et al., 1997; Tassoni et al., 2005). The tyrosine phosphatase inhibitor, sodium orthovanadate, is another known inducer of *VvSTS* (Tassoni et al., 2005).

Several R2R3-MYB and WRKY transcription factors (TFs) have been identified as STS regulators in different grapevine species in response to stress or developmental cues (Höll et al., 2013; Jiang et al., 2019; Vannozzi et al., 2018; Wang et al., 2019; Wong et al., 2016). Some of these regulators have been shown to act as activators and some as repressors of STS gene expression.

Protein synthesis inhibitors like cycloheximide (CHX) provide widely used and useful tools for identifying primary response genes (PRG) that do not require *de novo* protein synthesis but are regulated by pre-existing TFs and labile repressors. Conversely, secondary response genes (SRG) are dependent on the synthesis of new TFs after the stimulus and cannot be activated if the translation is inhibited (Tullai et al., 2007; Bahrami and Drabløs, 2016). It has been noticed that CHX treatments cause side effects such as inducing ROS production (Tenhaken and Rübel, 1998) and disturbing phosphorylation status by activating protein kinases, which need to be considered when interpreting the results (Usami et al., 1995).

In this study, the transcriptional response to UV-C treatment of pine needles was analyzed to discover regulators and missing enzymes of the Scots pine stilbene biosynthetic pathway. Such large-scale transcriptional analysis of UV-C responses has not been done in gymnosperm species before. Further, this study explored the induction of gene expression in the stilbene pathway during CHX treatment, which has not been studied in grapevine. In addition, we showed that the stilbene pathway is activated in pine in response to phosphatase inhibitors and plant hormones, as it is in grapevine. Pine STS promoter was able to activate the expression of a reporter gene in Arabidopsis, species that does not naturally produce stilbenes, in response to UV-C exposure and phosphatase inhibitors, but not as a response to plant hormone treatments. Several TFs and two 4CL encoding genes were coexpressed with stilbene synthase. However, their involvement in the stilbene pathway needs to be studied in detail with additional experiments.

## Results

### Induction of the stilbene pathway as a response to UV-C and CHX treatments

Six-week-old Scots pine seedlings were treated with UV-C and expression of the transcript encoding the key enzyme of the stilbene pathway, *PsSTS*, was followed for 48 hours with semi-quantitative RT-PCR (Fig 1A). Induction of *PsSTS* was evident four hours after the treatment and reached the highest level at 24 hours and then declined, showing lower expression level at 48 hours. Pinosylvin was detected from the needles 24 hours after the treatment (Fig S1).

**Figure 1.**
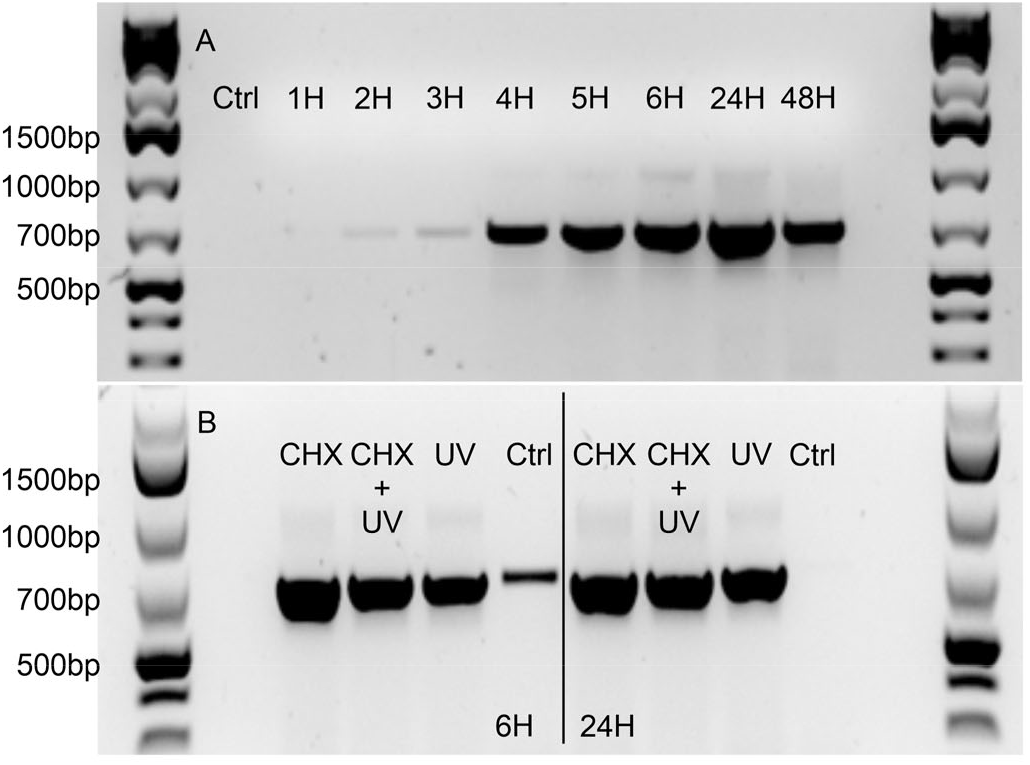
A) The expression of *PsSTS* was followed with RT-PCR for 48 hours after the onset of UV-C treatment. Expression of *PsSTS* was clearly visible after four hours of induction, strongest at 24 hours and started to decline after that showing lower expression level at 48 hours. B) The expression of *PsSTS* was analyzed six and 24 hours after onset of UV-C treatment. Plants were treated overnight with 10 μM of CHX prior the UV exposure. Expected size of the *PsSTS* fragment is 700 bp.

Since expression of *PsSTS* was induced relatively fast after the UV-C exposure, we tested if the expression would be independent of *de novo* protein synthesis. Translation was inhibited using an overnight treatment of CHX prior to the UV-C treatment. Inhibition of protein synthesis did not prevent the accumulation of *PsSTS* transcripts, and in fact, CHX treatment alone was able to activate *STS* gene expression (Fig1B). Based on the incorporation rate of radioactive methionine in needle proteins, CHX treatment inhibited translation by 94%. Another protein synthesis inhibitor, anisomycin, was also able to activate *PsSTS* gene expression (Fig S2).

### Transcriptional changes in response to UV-C treatment

Transcriptomes of needles from 6-week-old seedlings treated with UV-C and sampled from four Scots pine families at two, six and 24 hours after onset of the treatment were sequenced using SOLiD platform and compared to untreated control (Fig S3). In response to UV-C treatment, 6131 TCs (tentative consensus sequences) in the Pinus EST collection (The Gene Index Database, 2014) showed differential expression (false discovery rate, FDR < 0.05) in at least one of the time points when compared to control samples, based on analysis by edgeR (Table S1). Mapping was done against the EST collection instead of a Trinity assembly of our own data to observe the expression of genes that were not present in our data, as described in our previous studies (Lim et al. 2016, 2021).

To get a better view of the response to UV-C treatment, hierarchical clustering was used to capture specific expression patterns of transcripts. Sequence data was mapped against Pinus EST collection and 12709 transcripts that achieved at least eight counts per million (CPM) in at least four libraries were subjected to *SplineCluster* analysis, which uses Bayesian hierarchical clustering algorithm. Transcripts were grouped into 18 clusters according to their expression profiles (Fig 2, Table S2). Based on the analysis, four different expression patterns could be distinguished. In the first pattern, in clusters 1-7, the expression peaked either two or six hours after the treatment and then declined and in some cases returned to control level. In the second pattern, in clusters 8-14, transcription was activated after the treatment, but the peak of the expression was beyond the time points measured. In the third pattern, clusters 15-16, expression of transcripts were downregulated. Cluster 18 formed the final pattern where the expression pattern was flat. Each pattern was targeted to Gene Ontology (GO) enrichment analysis and the GO categories, biological processes, were the most informative (Fig S4A). In the upregulated patterns 1 and 2, GO terms such as response to stress, stimulus and signaling were enriched. Downregulated pattern 3 was enriched with photosynthesis-related transcripts.

**Figure 2.**
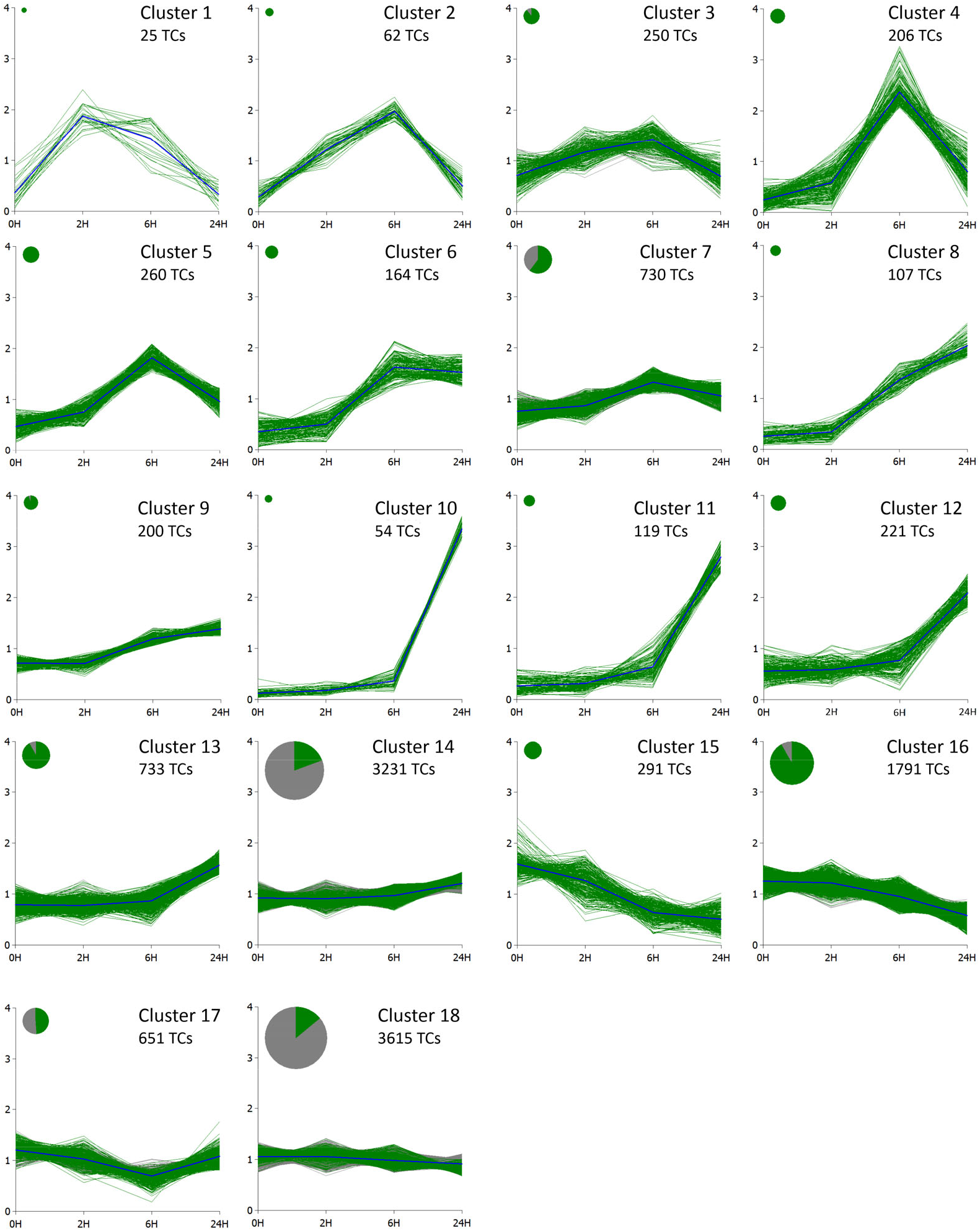
Transcripts from UV-C treatment data were grouped by the Bayesian hierarchical clustering algorithm *SplineCluster* (Heard et al., 2006) All TCs that obtained at least eight CPM in at least four libraries are shown. The transcripts that have statistically significant (FDR < 0.05) difference in expression levels compared to the control are shown in green. The area of the disc is proportional to the number of TCs in the cluster. 0H, control; 2H, 2 hours; 6H, six hours; 24H, 24 hours.

### Transcriptional responses of secondary metabolite pathways to UV-C stress

The expression of stilbene pathway encoding genes were upregulated in response to UV-C treatment. The transcripts were found in different clusters. Full length PsSTS and PsPMT2 enzymes are encoded by TCs TC154538 and TC166778, respectively (Schanz et al., 1992, Paasela et al. 2017). Both were found in cluster 8 where genes were upregulated but did not reach the peak expression in 24 hours. Most of the stilbene synthase transcripts in the Pinus EST collection followed the same pattern but some were found in cluster 6 where expression levels off during the time course. Scots pine has five stilbene synthase encoding genes (Preisig-Müller et al.,1999) but their coding sequence is so similar that the resolution of short SOLiD sequencing cannot tell them apart. Most of the PAL and 4CL (4-coumarate ligase) encoding transcripts followed the first pattern where the expression started to decline after the initial induction. They were present in clusters (3-7). Some could also be found in clusters 13 and 14 that showed mild up-regulation without decline in the expression.

The mevalonate (MVA) pathway genes, such as 3-hydroxy-3-methylglutaryl coenzyme A reductase (HMGR) were strongly upregulated by UV-C exposure and were present in clusters 3, 4 and 6. The MVA pathway is important for biosynthesis of essential terpenoid products such as sterols, sesquiterpenes, triterpenes, cytokinins and brassinosteroids (Rodríguez-Concepción & Boronat 2002). The sesquiterpene biosynthetic enzyme E,E-α-farnesene synthase was strongly upregulated in cluster 4. In contrast, genes encoding enzymes for the core chloroplastic isoprenoid producing pathway, the methylerythritol 4-phosphate (MEP) pathway, and enzymes downstream of the MEP pathway involved in monoterpene and diterpene resin acid biosynthesis were downregulated.

Lignin biosynthesis pathway genes cinnamoyl-CoA reductase (CCR), caffeoyl-CoA *O*-methyltransferase (CCoAOMT) and hydroxycinnamoyl-CoA:shikimate hydroxycinnamoyltransferase (HCT) were induced in response to UV-C treatment moderately, two to five fold and followed first expression pattern.

The expression of chalcone synthase, the key enzyme of flavonoid pathway, or other enzyme encoding genes in the pathway such as flavanone 3-hydroxylase, flavonoid 3’-hydroxylase and flavonoid 3’,5’-hydroxylase were not induced strongly by UV-C treatment. After 24 hours only some transcripts reached up to 2-fold upregulation.

All the TCs encoding the major secondary metabolite biosynthesis genes and their expression levels in response to UV-C treatment were collected in Table S3.

### Transcriptional changes in response to CHX treatment

CHX induces *PsSTS* expression (Fig 1B) and in order to choose informative time points for RNA sequencing, we performed a time series experiment to see how fast CHX was inducing the expression. Increased expression was visible already at two hours after applying CHX, but strong activation was seen in both replicates after eight hours (Fig S5). RNA sequencing was performed on samples treated with CHX for two, eight and 24 hours in addition to the untreated control. In response to the inhibition of protein synthesis by CHX, 9851 genes were differentially expressed compared to the untreated control (Table S4).

*SplineCluster* analysis was also done to CHX data. After mapping against the Pinus EST collection, 18239 transcripts achieved at least eight CPM in at least four libraries and were kept. These were subjected to hierarchical clustering and were grouped into 19 clusters according to their expression profiles (Fig 3, Table S5). Responses to CHX can be divided into similar expression patterns as UV-C data. Clusters 1-4 followed the first pattern where the expression peaked eight hours after the treatment and then declined and, in some cases, returned to the control level. In the second pattern, in clusters 5-13 transcription was activated after the treatment, but the peak of the expression was not seen within the time points measured. In the third pattern clusters 14-17 expression of transcripts were downregulated. The fourth expression pattern was flat including clusters 18-19. In total 36% of transcripts deposited in *Spline cluster* analysis can be considered as primary response genes based on the definition and presence in clusters 1-13.

**Figure 3.**
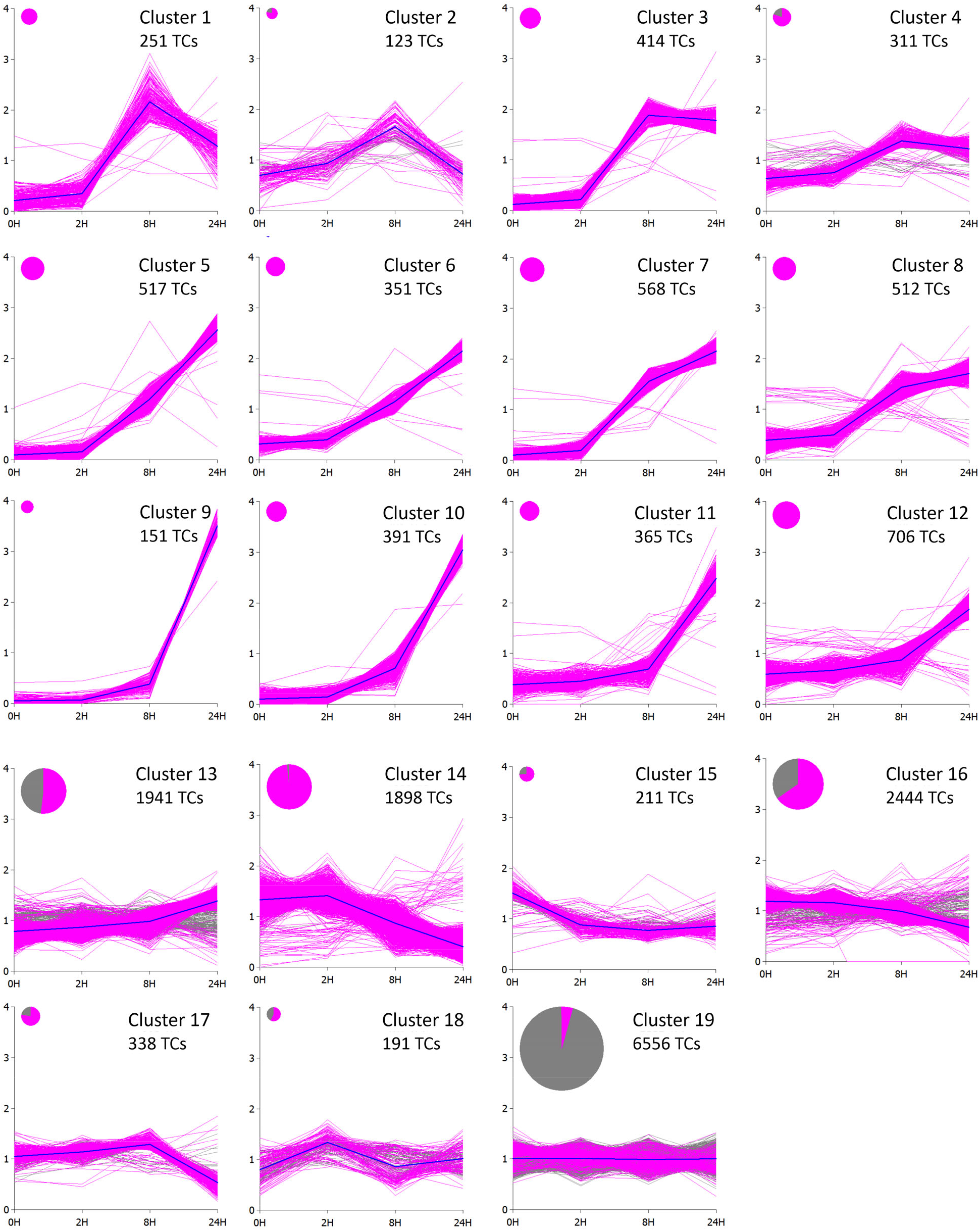
Transcripts from CHX treatment data were grouped by the Bayesian hierarchical clustering algorithm *SplineCluster* (Heard et al., 2006) All TCs that obtained at least eight CPM in at least four libraries are shown. The transcripts that have statistically significant (FDR < 0.05) difference in expression levels compared to the control are shown in pink. The area of the disc is proportional to the number of TCs in the cluster. 0H, control; 2H, 2 hours; 8H, eight hours; 24H, 24 hours.

GO enrichment analysis of CHX treated seedlings showed similar GO terms as UV-C treated. (Fig S4B). The upregulated patterns 1 and 2 have transcripts involved in signaling, stress and jasmonic acid responses. Downregulated pattern 3 has GO terms related to photosynthesis.

### Transcriptional responses of secondary metabolite pathways to CHX treatment

Different secondary metabolism pathways responded differently to CHX treatment. Full length *PsSTS* and *PsPMT2* genes were found in cluster 9 following the second pattern and were upregulated but did not reach the peak of expression during the measured time points. Additional transcripts were present also in cluster 10, following the same pattern. PAL and 4CL encoding transcripts were found both in inducible clusters following pattern two: 5-7, 10,11,13 and 14, downregulated clusters 14 and 16 and non-inducible cluster 19 of CHX treatment.

The expression of chalcone synthase was downregulated in response to CHX treatment, and the TCs were mostly in clusters 14 and 16. Plastidic terpenoid biosynthesis encoding genes such as the core MEP pathway and downstream genes involved in mono- and diterpene biosynthesis were downregulated. MVA pathway, in contrast, was strongly upregulated by CHX treatment. One specific sesquiterpene synthase encoding gene strongly upregulated in response CHX was E,E-α-farnesene synthase, present in cluster 5. Several dirigent proteins involved in lignan biosynthesis were upregulated in CHX data and were present in cluster 5. Genes specific for the lignin pathway were slightly upregulated in clusters 12 and 13 or downregulated in cluster 14. Expression data of secondary metabolite pathway encoding genes in response to CHX treatment were collected in Table S3.

### Novel candidates for uncharacterized enzymes of pine stilbene pathway

The specific PsPAL and Ps4CL enzymes are not characterized in the Scots pine stilbene pathway. The *PsSTS* and *PsPMT2* encoding genes (TC154538 and TC166778, respectively) were present in the same clusters both after UV-C and CHX treatments. These clusters, however, did not contain any PAL or 4CL encoding genes. Several PAL and 4CL encoding TCs were present both in other upregulated and downregulated clusters after both UV-C and CHX treatments. What makes their analysis complicated is that these enzymes are shared with stilbene, lignin, lignan and flavonoid pathways. Different secondary metabolite pathways seemed to respond differently at the transcriptional level to the treatments studied here when specific enzymes of each of the pathways are compared. Stilbene synthase was strongly upregulated, chalcone synthase was downregulated or remaining constant and HCT and CCR from lignin pathway were slightly upregulated. This difference is especially clear in the CHX data. All transcripts encoding PAL and 4CL present in the CHX data were collected and plotted to visualize different expression patterns (Fig S6). The best candidates were TC172794 encoding PAL which was highly upregulated in response to CHX treatment and two 4CL encoding TCs TC179200 and TC191762 that were induced at similar levels as the characterized PsPMT2 enzyme. Simple blast of known CNLs amino acids, enzymes capable of activating cinnamate in petunia and Aaron’s beard, to pine TC collection did not reveal any similar sequences.

Expression values of the core stilbene biosynthetic pathway enzyme encoding genes were collected in Table S3.

### Novel transcription factor candidates for the pine stilbene pathway

No regulators of the stilbene pathway in pine have been characterized. To predict what kind of transcription factors might be involved in regulating the stilbene pathway we did an *in silico* promoter analysis of the Scots pine *STS* promoter PST-1 (Preisig-Müller et al.,1999) using PlantPan 2.0 (Chow et al., 2016) software to screen putative TF binding elements. According to Preisig-Müller et al.

(1999), the PST-1 promoter has the highest responsiveness of all the *PsSTS* promoters when tested in transgenic tobacco. The promoter contains one or several binding sites for several TF families such as NAC, MYB, WRKY, bZIP, bHLH, AP2/ERF, Homeodomain and GRAS (Supplemental Table S6).

To find putative regulators of the Scots pine stilbene pathway all differentially expressed transcripts associated with GO terms GO:0003700 (sequence-specific DNA binding transcription factor activity) or GO:0003677 (DNA binding) were extracted from UV-C and CHX RNA-sequencing data and from previously published transition zone/sapwood (TZ/SW) (Lim et al. 2016) and wounding data (Lim et al. 2021), two other conditions where stilbene pathway is activated. TFs from all these different treatments and tissues were compared. A Venn diagram was built from the extracted transcription factor data to identify common regulators (Fig 4A) Only four genes encoding transcriptional regulators were common in all conditions, two WRKYs, one ERF and one IAA protein. Two additional TFs, ccaat-box binding factor hap3 and nuclear transcription factor y subunit b-, can be added to the list if we leave CHX treatment out from the comparison (Table S7). None of the MYB TFs was found to be commonly expressed.

**Figure 4.**
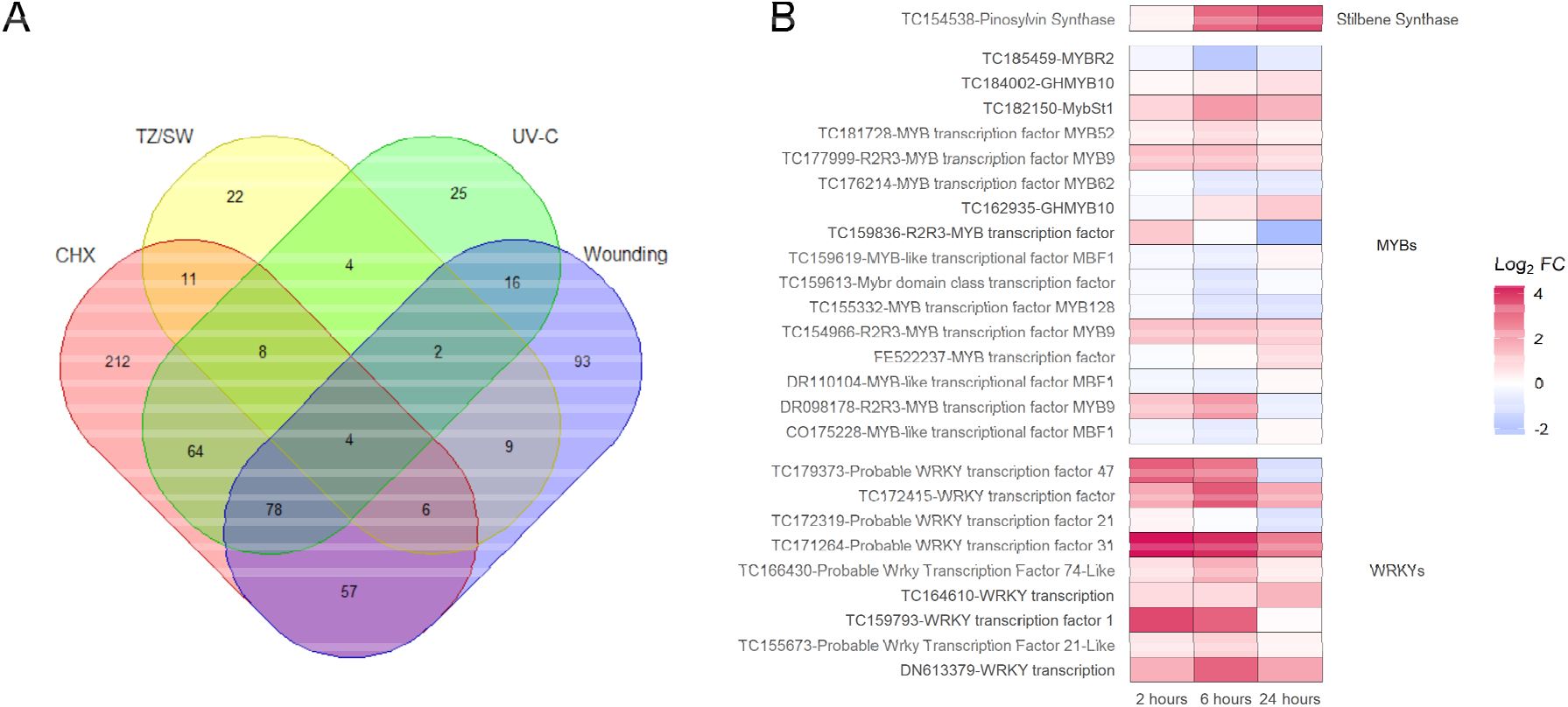
All differentially expressed transcripts associated with GO terms GO:0003700 (sequence-specific DNA binding transcription factor activity) or GO:0003677 (DNA binding) were extracted from UV-C and CHX RNA-seq data and from previously published transition zone/sapwood (TZ/SW) (Lim et al. 2016) and wounding data (Lim et al. 2021. A Venn diagram was built from the extracted transcription factor data to see the common regulators in each condition grouping (A). Expression of MYB and WRKY TFs, the most likely candidates for *PsSTS* transcriptional regulation, from UV-C data were further compared to *PsSTS* expression to reveal co-expressed TFs.

It is possible that the stilbene pathway in Scots pine is regulated via different TFs in different inducible conditions. Since R2R3-MYB and WRKY TFs regulate *VvSTS* expression, all differentially expressed TFs from these families were extracted from UV-C data and their expression was compared to *PsSTS* expression (Fig 4B). Three R2R3-MYB (TC177999, TC154966 and DR098178) and four other types of MYB (TC184004, TC182150, TC181728 and TC162935) TFs showed inducible expression patterns in response to UV-C treatment. All the WRKYs showed some level of induction as a response to UV-C treatment. The closest ortholog for grapevine R2R3-MYB14 and 15 and *Picea glauca* MYB12 (ABQ51228) (Höll et al., 2013) in pine is R2R3-MYB TC182032. This TC was strongly expressed in CHX data and showed an inducible expression pattern in response to UV-C treatment but was filtered out because it did not fulfill the cutoff criterion of eight CPM in at least four libraries. The candidate MYB TF TC188897 that was co-expressed with *STS* in the TZ (Lim et al. 2016) was strongly activated in response to CHX treatment but not in response to wounding or UV-C. The NAC TF (TC164798) that was expressed in the TZ and following expression of pine stilbene pathway enzymes (Lim et al. 2016) was not differentially expressed in any other treatment in addition to TZ. The expression profiles of commonly expressed TFs and MYB candidates that were not present in UV-C data R2R3-MYB TC182032 and MYB TC188897 are shown in Table S7.

### Induction of stilbene synthase expression in response to phosphatase inhibitors and plant hormones ethylene and jasmonate

We wanted to test if the other known inducers of the stilbene pathway in grapevine, the tyrosine phosphatase inhibitor, sodium orthovanadate (Tassoni et al., 2005) and plant hormones jasmonate and ethylene would induce the pathway also in pine (Belhadj et al, 2008; Grimmig et al., 1997; Tassoni et al., 2005). First, the effect of inhibition of protein phosphatases was tested with a broad-spectrum inhibitor cocktail (Fig S7). The treatment strongly activated the expression of *PsSTS* with similar kinetics as CHX and UV-C. Next, the inhibiting effect of protein phosphatases were tested using more specific inhibitors targeting tyrosine phosphatases separately with sodium orthovanadate and Ser/Thr phosphatases with sodium pyrophosphate (Fig 5) and β-glycerophosphate (Fig S8). Combined effects were tested as well (Fig S8). Tyrosine phosphatase inhibitor had a much stronger activating effect on the expression of the stilbene synthase gene than Ser/Thr phosphatase inhibitors did. Ser/Thr phosphatase inhibitors had only a mild effect on *PsSTS* gene expression.

**Figure 5.**
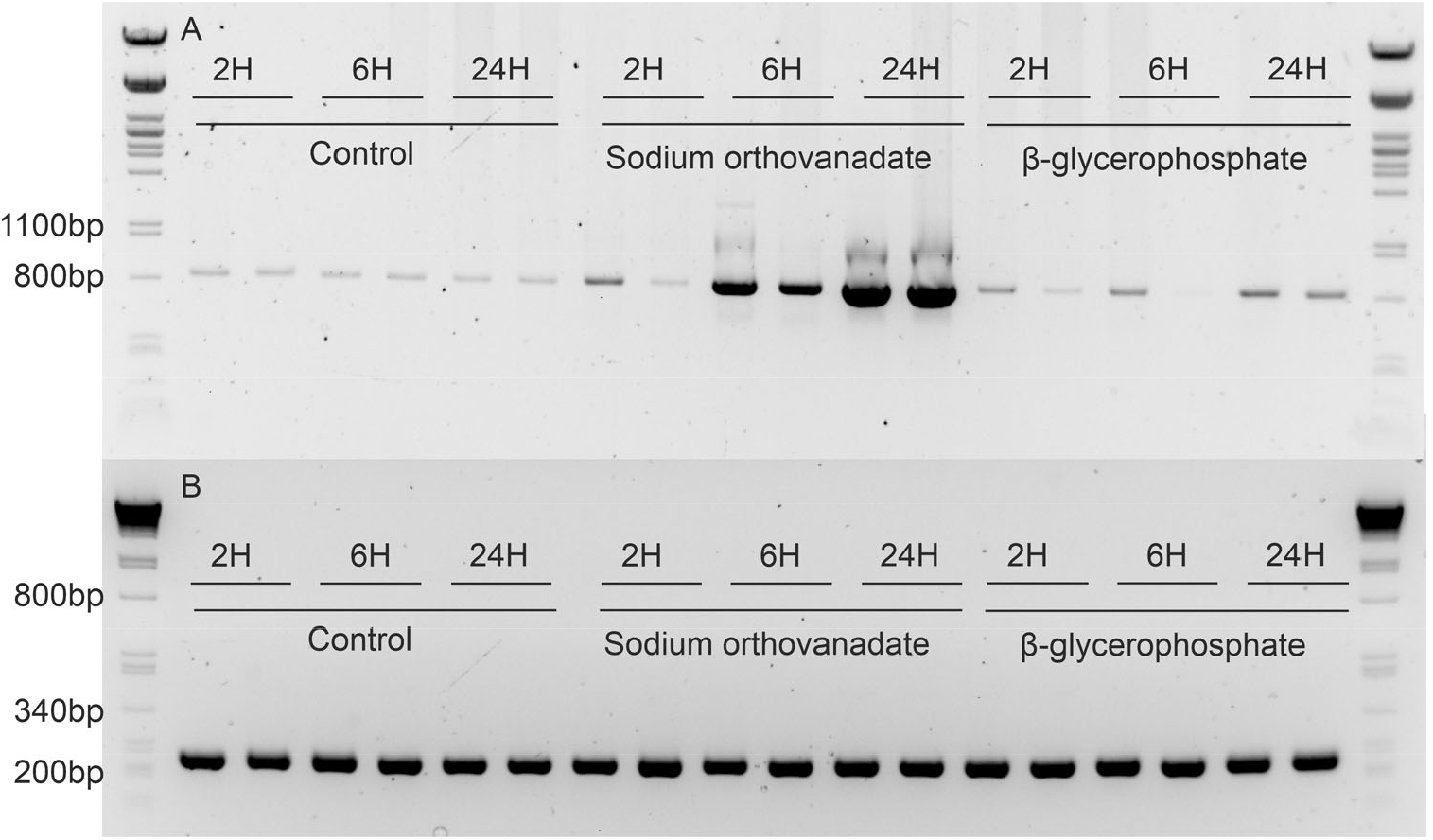
A) The expression of *STS* in pine needles was analyzed after two, six and 24 hours of treatment with 1mM sodium orthovanadate, 1mM β-glycerophosphate or from untreated control plants. Strong induction of *STS* gene expression was seen six hours after the treatment. Slight induction of *STS* was seen 24H of β-glycerophosphate treatment. Expected size of the *PsSTS* fragment is 853 bp. B) The expression of housekeeping gene *actin* was used as a control.

We tested the response of the key enzyme of stilbene pathway, *PsSTS*, to ethylene and JA. Ethylene and JA did not cause consistent increase of *PsSTS* expression between different experiments and replicates when applied separately, but when applied simultaneously, the induction was strong and repeatable (Fig 6).

**Figure 6.**
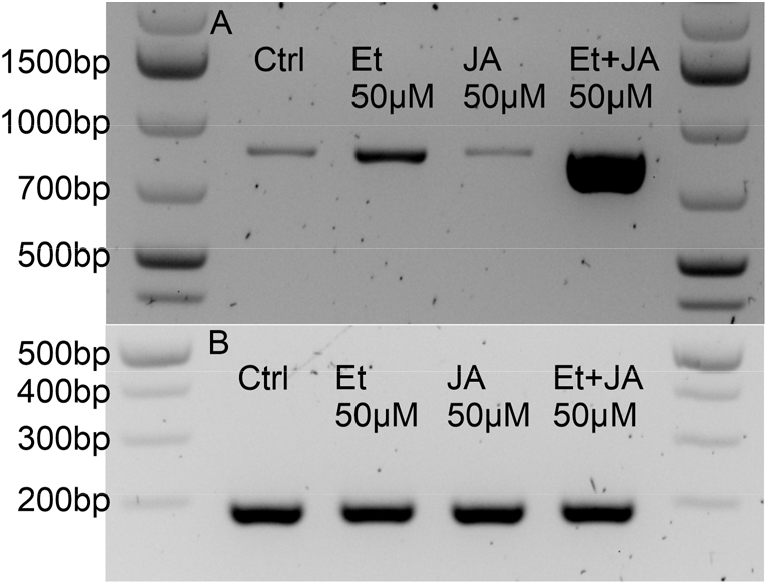
A) Induction of *PsSTS* in response to treatment with 50 μM ethylene (Et) and 50 μM jasmonic acid (JA) or in the combination of both (50 μM Et+50 μM JA). Ethylene and jasmonate separately caused erratic activation of *PsSTS* but when applied together induction was strong and consistent. Expected size of the *PsSTS* fragment is 853 bp B) Gene encoding *actin* was used as a control.

### Regulation of the pine stilbene synthase promoter PST-1 in non-stilbene producing plant

Scots pine has five genes encoding stilbene synthase, which were named PST-1 to PST-5 (Preisig-Müller et al. 1999). It was shown that pine stilbene promoters activate reporter gene expression in response to UV-C, wounding and pathogen infection in species that do not normally produce stilbenes such as in tobacco (*Nicotiana tabacum*) where PST-1 showed the highest responsiveness (Preisig-Müller et al. 1999). We transformed Arabidopsis with the PST-1 promoter combined with the firefly luciferase (LUC) reporter gene. We tested if the PST-1 promoter can drive LUC gene expression in response to the same treatments found to induce *PsSTS* gene expression in pine. Clear luciferase activity was detected in plants that were exposed to UV-C or treated with phosphatase inhibitor. However, the pine promoter was not able to activate luciferase expression in response to combined treatment with plant hormones ethylene and JA (Fig 7).

**Figure 7.**
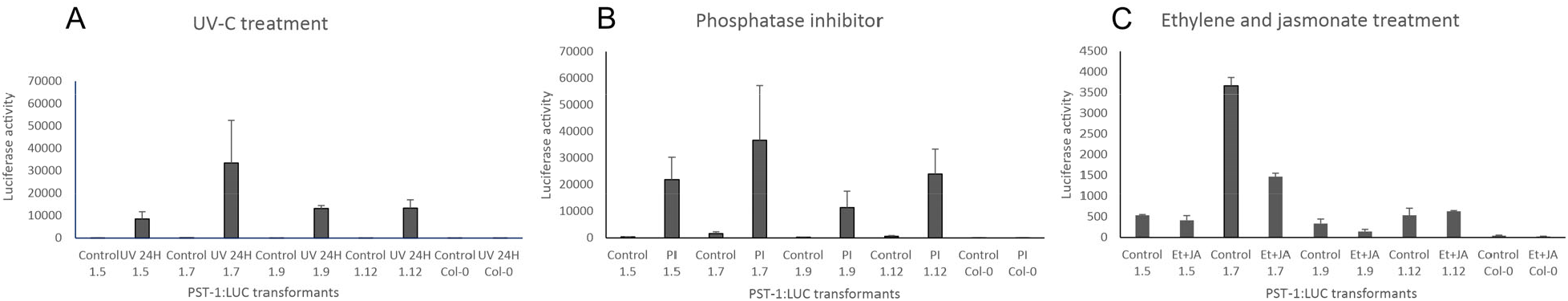
Four individual transformants (1.5, 1.7, 1.9 and 1.12) of Arabidopsis carrying PST-1:LUC promoter-reporter construct were treated with UV-C (A), phosphatase inhibitor cocktail (B) or combination of 50 μM Ethylene precursor ACC and 50 μM of jasmonate (C). Plants were sampled before and after the treatment and LUC activity was measured. UV-C and phosphatase inhibitor treatment resulted in increased luciferase activity in all the independent transformants. Hormonal treatment had no effect on LUC activity. Error bars represent standard error from 3-5 replicates.

## Discussion

UV-C treatment was the first discovered artificial stress inducer of the stilbene pathway in pine (Schoeppner & Kindl 1979). Large-scale analysis of transcriptomic responses to UV-C has been utilized in grapevine to study plant responses to the treatment and to discover regulators of the pathway (Höll et al. 2013, Vannozzi et al. 2012, Xi et al. 2014). In this study we exposed 6-week-old pine seedlings to UV-C treatment and analyzed their needle transcriptomes. Our main aim was to discover co-expressing TFs as candidates for regulating the stilbene pathway in pine and find the pathway-specific cinnamate activating CoA ligase. In general, UV-C caused massive changes in gene expression in pine with 6131 differentially expressed TCs in at least one of the analyzed time points. The GO enrichment analysis showed similar responses in pine as is seen in grapevine. In brief, stress response pathways were upregulated and chloroplast and photosynthesis genes were downregulated.

Protein synthesis inhibitors like CHX, are used for identifying whether the initiation of gene expression requires *de novo* protein synthesis or is regulated by pre-existing TFs (Tullai et al., 2007; Bahrami and Drabløs, 2016). The definition of a primary response gene is that its transcription is initiated without de novo synthesis of transcriptional regulators. The kinetics of the initiation of transcription vary among the primary response genes and they can be further categorized as primary response genes and delayed primary response genes (Tullai et al., 2007). Treating plants with translational inhibitor CHX caused even bigger changes in gene expression than UV-C treatment did. In total 9851 genes were differentially expressed compared to the untreated control. In total 36% of transcripts deposited in the *SplineCluster* analysis can be considered as primary response genes based on the definition and presence in clusters 1-13 in which the transcription was activated after the treatment. However, we cannot completely rule out the possibility that some of the responses we see for CHX treatment are due to secondary effects of the treatment such as CHX induced ROS production (Tenhaken and Rübel, 1998) or disturbance of the phosphorylation status (Usami et al., 1995) instead of translational inhibition *per se*. Based on GO analysis, after CHX treatment upregulated clusters had biological processes such as signaling and stress enriched and in downregulated clusters photosynthesis related terms were enriched.

### Phenylpropanoid pathways respond to UV-C exposure and CHX treatment in different ways

In pine and grapevine, UV-C exposure causes strong transcriptional activation of the stilbene pathway and subsequent accumulation of stilbenes (Vannozzi et al., 2012; Xi et al., 2014; Suzuki et al., 2015). Transcriptional activation of the stilbene pathway in needles was clearly seen in the RNA-seq data and the product pinosylvin was detected 24 hours after the treatment. Several pinosylvin synthase encoding TCs and an enzyme encoding gene downstream of the pathway, *PsPMT2*, were upregulated. In grapevine, induction of the stilbene pathway suppressed the expression of CHS and biosynthesis of flavonoids (Jeandet et al., 1995; Vannozzi et al., 2012). It has been shown that in tissues and developmental stages where the flavonoid pathway is active, STS expression is very low (Jeandet et al., 1995; Vannozzi et al., 2012). Furthermore, it has been shown that in *Pinus densiflora* pinosylvin inhibits CHS enzyme activity (Kodan et al., 2002). We saw that the lignin pathway-specific enzymes HCT and CCR were only slightly induced by the UV-C, but the flavonoid pathway encoding genes, such as *CHS* did not show induction. It is still possible that stilbene pathway is induced at the expense of flavonoid pathway because they use the same precursors, and this feature seems to be conserved between angiosperms and gymnosperms.

For the pine stilbene biosynthetic pathway, the genes encoding pathway specific PAL and cinnamate activating enzymes are not characterized. In Aaron’s beard and petunia cinnamate is activated by cinnamate:CoA ligase (CNL) in the biosynthetic pathway of benzoic acid and its derivatives (Klempien et al. 2012, Gaid et al. 2012). We could not find genes encoding CNL-like enzymes from the Pinus EST collection. Cinnamate can also be converted by a specific 4CL variant like in raspberry (Kumar & Ellis 2003). Several PAL and 4CL encoding TCs were present both in upregulated and downregulated clusters in both UV-C and CHX treatment transcriptomes. From the phenylpropanoid pathways only stilbene pathway genes seem to react strongly to CHX treatment, and this might help to find enzymes upstream of STS. From the transcriptomic data one inducible PAL was discovered. TC172794 encoding PAL was highly upregulated in response to CHX treatment. Two 4CL encoding TCs TC179200 and TC191762 were induced at similar levels as the gene for the PsPMT2 enzyme. The capability of these enzymes to activate cinnamate needs to be shown in additional experiments.

### Regulators of pine stilbene pathway

Since the stilbene pathway in grapevine, and phenylpropanoid and flavonoid pathways in many species are regulated by MYB family TFs, it is the most likely candidate family to regulate the pine stilbene pathway as well (Bomal et al., 2008, 2013; Falcone Ferreyra et al., 2012; Höll et al., 2013; Patzlaff et al., 2003; Wong et al., 2016). In addition, recent data shows that WRKY TFs are coregulating (Vannozzi et al., 2018; Wang et al., 2019) or inhibiting (Jiang et al., 2019) *VvSTS* expression and AP2/ERF are present in the stilbene pathway regulatory network in grapevine (Wong et al., 2016). It has been shown that many defense genes are regulated by the synergistic action of ethylene and jasmonate signaling pathways, and in many cases the TF responsible for the regulation is ethylene response factor 1 (ERF1) (Lorenzo et al., 2003). The same TFs, VvMYB14 and VvMYB15, mediate *VvSTS* gene induction and stilbene production both in stress induction when plants were wounded, infected with downy mildew, or treated with UV-C, and based on developmental cues (Höll et al., 2013). However, different regions of the STS gene promoters are responsible for pathogen and ozone responsiveness, and ozone and ethylene activate *VvSTS* expression via separate signaling pathways (Grimmig et al., 1997; Schubert et al., 1997).

The expression patterns and timing of different members of the large Vitis *STS* gene family vary in response to different stress treatments, which indicates transcriptional subfunctionalization in responsiveness to different signaling pathways (Vannozzi et al., 2012). Scots pine stilbene synthase is encoded by a family of five genes (Preisig-Müller et al.,1999), but we do not yet know if there is a division of labor between the different family members in different conditions such as environmentally and developmentally induced production as it is in grapevine. With short RNA-seq reads it is difficult to tell apart the different paralogs. We know from previous studies that stilbene biosynthesis in response to wounding stress and developmental induction is linked in some way. The offspring of individuals having the highest amount of pinosylvin in their heartwood, produce more stilbenes as a response to wounding of xylem (Harju et al., 2009).

The specific TFs regulating the stilbene pathway in pine are not known. *In silico* promoter analysis of *PsSTS* revealed several putative TF binding elements in the promoter of pine stilbene synthase encoding gene. In our previous study, two TFs belonging to the MYB and NAC families were shown to co-express with STS in pine in developmental context (Lim et al., 2016), but these TFs were not expressed in the UV-C libraries, which might indicate that different factors are regulating developmental and stress responses or these TFs are not regulating *PsSTS* at all. A different set of TF encoding genes from different families were induced in response to UV-C and several good candidates were obtained from the experiment. Particularly interesting candidates are the VvMYB 14 and 15 homolog R2R3-MYB (TC182032), and WRKY (TC166430) and ERF (TC157450) TFs expressed in all stilbene producing conditions. However, additional experiments are needed to show which ones, if any, are involved in the regulation of the pine stilbene pathway.

A further complication for finding the regulators of pine stilbene pathway is brought by the fact that CHX treatment induces the expression of stilbene pathway genes, which suggests that pre-existing transcriptional machinery would already be present in the cells and simply activated in response to the signal. This is a common means to regulate, for instance hormone responsive genes, where activating TFs are counteracted by labile repressors. Under CHX treatment, labile repressors cannot be replaced after their degradation and transcription is activated (Moore et al., 2011). In grapevine, VvWRKY8 represses stilbene synthase gene expression by direct interaction with VvMYB14 and this way preventing its binding to the promoter (Jiang et al., 2019). We showed that the stilbene pathway genes (*PsSTS* and *PsPMT2*) were transcriptionally activated with slower kinetics than most of the primary response genes and resembled this way more delayed primary response genes (Tullai et al., 2007). On the other hand, some genes such as JAZ repressors are transcriptionally activated both in response to CHX and treatments such as wounding (Chung et al., 2008), which supports using transcriptional data for finding candidate regulators. So far, there is no experimental evidence of CHX induction of stilbene pathway genes in other stilbene producing species such as grapevine.

### Many aspects of stilbene pathway regulation are shared between angiosperms and gymnosperms

We know from previous studies that the stilbene pathway is activated in pine, for instance, in response to UV-C, wounding, fungal infection and ozone fumigation (Schoeppner&Kindl 1979, Harju et al., 2009, Lange et al. 1994, Zinser et al., 2000). However, the mechanisms and components of the signaling pathways under different inductive conditions of pine stilbene biosynthesis are still unknown.

It has been suggested that the plant hormone ethylene regulates the biosynthesis of stilbenes also in Scots pine. Treatment of pine sapwood with ethylene was shown to induce the production of pinosylvin and its monomethyl ether (Nilsson et al., 2002). The ethylene biosynthesis gene encoding 1-aminocyclopropane-1-carboxylate oxidase (ACO) was induced in the TZ where the accumulation of stilbenes and conversion of sapwood to heartwood occurs. However, the year around expression profile of *ACO* did not support the involvement of ethylene in the stilbene pathway induction (Lim et al., 2016). In this study, it was shown that treatment of pine seedlings with the ethylene precursor 1-aminocyclopropane-1-carboxylic acid (ACC) caused only erratic induction of *STS* expression in needles. JA treatment did not induce the *STS* gene expression either, but combination of these two hormones caused strong and reproducible activation of *PsSTS*. Defense genes such as chitinase (Lorenzo et al., 2003), plant defensin gene *PDF1*.*2* (Penninckx et al., 1998), pathogenesis related proteins *PR-1b* and *PR5* (Xu et al., 1994) and proteinase inhibitor (O’Donnell et al., 1996) are examples of defense and wound responsive genes where both hormones are needed for activation of the gene expression. It is possible that JA and ethylene regulate the expression of stilbene synthase gene in stress induced conditions but not in the developmental context.

In grapevine sodium orthovanadate, inhibitor of tyrosine phosphatase, induced the expression of stilbene synthase genes (Tassoni et al. 2005). This is another conserved feature in regulation of stilbene pathway between grapevine and pine. When we disturbed the phosphorylation status of Scots pine cells with protein phosphatase inhibitors, expression of stilbene synthase was activated, suggesting that at some level the stilbene pathway is under negative regulation of protein phosphatases. Sodium orthovanadate as tyrosine phosphatase inhibitor had a stronger effect on activation of the transcription than the tested Ser/Thr phosphatase inhibitors. Phosphorylation and dephosphorylation of TFs have been shown to be important mechanisms for initiating the transcription of PRGs. The TFs regulating PRG can be both activated and deactivated by phosphorylation. (Moore et al., 2011; Bahrami and Drabløs, 2016).

## Conclusion

It has been previously reported that pines synthesize stilbenes in response to wounding, ozone fumigation, UV-C exposure, pathogen attack and during heartwood development. In Scots pine, no large-scale transcriptomic studies have been done before in respect to UV-C treatment as has been done for grapevine. Activation of stilbene pathway but not of the flavonoid pathway occurs in response to UV-C both in grapevine and in Scots pine. New candidates for specific regulatory transcription factors co-expressed with stilbene synthase gene expression were identified, as well as genes encoding putative PAL and 4CL-like encoding genes specific for stilbene pathway. Furthermore, we showed that the pine stilbene pathway is regulated by protein (tyrosine) phosphatases and synergistic action of plant hormones ethylene and jasmonate, both influencing stilbene pathway expression in grapevine as well, except that in grapevine action of a single hormone, either ethylene or JA is enough for pathway activation.

Regulation of the Scots pine stilbene synthase gene expression is interesting. Even when the stilbene synthase gene has evolved from *CHS* (Tropf et al., 1994), it seems to be under different regulation. Stilbene synthase encoding genes are responding to CHX treatment showing features of fast responding primary responses, unlike CHS encoding genes. When the pine stilbene synthase gene promoter is transformed into plants that normally do not synthesize stilbenes, the stress inducibility is retained. This and shared features between gymnosperms and angiosperms regulation indicates that at least partly stilbene synthase regulation occurs via ancient stress-response pathway(s).

## Materials and methods

### Chemicals

All chemicals were purchased from Sigma-Aldrich unless stated otherwise.

### PST-1 promoter-LUC reporter vector construction and Arabidopsis transformation

DNA was isolated from pine needles using CTAB method as described in (Michiels et al. 2002). The PST-1 promoter was amplified from genomic DNA with primers attB4 PST-1-F and attB1r PST-1-R (Table S8). The PCR product was inserted into pDONRzeoP4P1r vector using Gateway BP clonase enzyme (ThermoFisher Scientific) according to the manufacturer’s instructions. Firefly luciferase gene with an intron (Mankin et al. 1997) was first amplified from plasmid pLKB10 with primers attB1 FLUC-F and attB5 FLUC-R and the PCR product was further amplified with primers Adapter attB1 and Adapter attB2 to generate the full attB1 and attB2 attachment sites. This PCR product was then inserted in pDONR221 vector. Using the Gateway BP reaction. Multisite Gateway cloning was done using LR clonase II Plus enzyme to fuse the PST-1 promoter fragment (attL4-attR1sites), the LUC fragment (attL1-attL2 sites) and a nopaline synthase terminator with attR2-attL3 sites to the pCAMkan-R4R3 destination vector. The final reporter construct PST-1:LUC:NOS was then transferred to Arabidopsis Columbia-0 ecotype using agrobacterium GV3101(pMP90) and the floral dip method (Clough & Bent 1998). Three rounds of selection were done in MS plates having 50 μg/ml of kanamycin. First primary transformants were selected. T1 lines with one insertion were selected by looking for 1:3 segregation. T2 lines lacking wild type segregants were propagated to establish homozygous T3 lines. Four independent transformants were used in all the experiments.

### Plant growth conditions

For UV-C and chemical treatments, pine seedlings were grown for six weeks and Arabidopsis for three weeks at 23 ºC in peat:vermiculite (1:1) under 16 h light, 8 h dark in controlled growth chambers. For the RNA sequencing experiments the Scots pine seeds originated from Natural Resources Institute progeny trial from Leppävirta (62°25′ N and 27°45′ E) and for the other experiments the seeds were a gift from Siemen Forelia and were collected from Hartola (61°36′ N and 26°16′ E).

### UV-C treatment

For the UV-C experiment, pine and Arabidopsis seedlings were treated for 15 min, at the distance of 20 cm, under an uncovered mercury lamp (sterilAir UVC G9). Spectral photon irradiance (μmol m^-2^ s^-1^) transmitted by the lamp was measured with an array spectroradiometer, which had been calibrated for measurements of UV and visible radiation (Maya2000 Pro Ocean Insight, Dunedin, FL, USA; D7-H-SMA cosine diffuser, Bentham Instruments Ltd, Reading, UK). The main peak from spectrum was at 254 nm (Fig S8).

The preliminary screen for pine UV-responses was done with single seedlings. Transcriptomes were sequenced from control samples and after two, six and 24 hours after onset of the UV-C treatment. Each sequenced sample had needles from ten seedlings pooled together, and each time point had four replicate samples (seedlings from open-pollinated mother tree lines T089, T170, T434 and T474). Samples were frozen in liquid nitrogen and stored at -80 °C.

For Arabidopsis, five plants from four independent transformants were treated with UV-C. Samples for luciferase measurements were taken before and after a 24 hours treatment. The effect of cutting of the leaf was tested by sampling the same non-treated plants 24 hours after first sampling (Fig S9A).

### CHX treatment

For all CHX treatments, seedlings were removed from the soil and the roots were rinsed with water. Roots were rolled around a pipette tip and seedlings were placed in 1.5 ml microcentrifuge tubes. For the preliminary screen (Fig 1), single plants were treated for 16 hours with 10 μM CHX in 1 ml of Milli Q (MQ) water in microcentrifuge tubes and the UV-C exposure was done the following day.

To test how efficiently protein translation was inhibited, ^35^S-methionine was fed to the plants through roots. 40 μl of 10 mCi/ml EasyTag™ Methionine, L-[35S] (Perkin-Elmer) was combined with 20 μl of MQ water. Seedlings were placed in microcentrifuge tubes and 10 μl of mixture was given to dry roots. Every hour 100 μl of MQ water was added to prevent seedlings from drying out. Treatment was done for four hours to two control plants and to four plants pre-treated with 10 μM or 100 μM of CHX for 16 hours. Samples were frozen in liquid nitrogen and stored at -80 °C. Proteins were extracted from the needles as described in Fliegmann et al. (1992) and were precipitated with ¼ volume of 100 % (w/v) trichloroacetic acid from part of the sample and the pellet washed twice with cold acetone. Radioactivity was measured from the extract and the trichloroacetic acid precipitated protein pellet for 5 min with a scintillation counter (Wallac 1414 WinSpectral v.1.40; Wallac Oy, Turku, Finland) using 1 ml of Ultima Gold™ (PerkinElmer) liquid scintillation cocktail. Labelling of newly formed polypeptides showed a reduction of 94% of protein synthesis in samples pre-treated with CHX compared to water control.

For CHX RNA-seq analysis ten 6-week-old seedlings were combined for each time point and placed in 10 ml of MQ water supplemented with commercial plant fertilizer (Substral, Scotts) and allowed to recover overnight. Next day the water was replaced with 10 ml of 10 μM CHX dissolved in MQ water or 10 ml of MQ water for the controls. Needles were collected two, four, six, eight, ten and 24 hours after initiation of the treatment. Control samples were collected at time points 2 and 24 hours. The following samples were sequenced from two biological replicates (lines T392 and T432) using SOLiD platform: control (24 hours in water), two hours after onset of the CHX treatment for early responses, eight hours when STS expression was clearly up in both replicates and 24 hour samples when the expression of STS was at its highest. Samples were frozen in liquid nitrogen and stored at - 80 °C.

### Other chemical treatments

For hormone, phosphatase inhibitor and anisomycin treatments, pine and Arabidopsis seedlings were removed from soil and placed in 1ml of MQ+Substral and allowed to recover overnight. The next day plants were treated with either 50 μM of ACC, 50 μM of JA or both combined, and samples were collected 6 hours and 24 hours after initiation of the treatments. The ACC stock was dissolved in water and JA in ethanol, and controls were treated with a similar volume of ethanol without hormones. Five plants were pooled together for each sample for the pine experiment. Samples were frozen in liquid nitrogen and stored at -80 °C. For Arabidopsis, five plants were treated from each line. Plants were sampled before and 24 hours after the treatment and luciferase activity was measured directly from fresh samples.

The broad-spectrum phosphatase inhibitor (PhosSTOP, Phosphatase Inhibitor Cocktail Tablet, Roche) was dissolved in MQ water and 1x solution was used for the treatments. Treatment was done for a single pine seedling per time point and five Arabidopsis plants per each line. The tyrosine phosphatase inhibitor sodium orthovanadate was prepared in alkaline water according to the manufacturer’s information. Ser/Thr inhibitors β-glycerophosphate and sodium pyrophosphate stocks were prepared in MQ water. Treatment was done with 1 mM of inhibitor solutions or alkaline MQ water, pH 10 for controls. Samples were frozen in liquid nitrogen and stored at -80 °C for pine samples and Arabidopsis plants were sampled before and 24 hours after the treatment and luciferase activity was measured directly from fresh samples.

For anisomycin treatment, inhibitor stock was dissolved in MQ. After the recovery period, MQ was replaced with 10 μM, 25 μM, 50 μM or 100 μM anisomycin and MQ water for the control plants. Samples were frozen in liquid nitrogen and stored at -80 °C.

### Extraction of total RNA and semiquantitative RT-PCR

Total RNA was extracted from needles as described in Chang et al. (1993) and dissolved in 50 μl of MQ water. RNA was purified and genomic DNA digested with RNeasy Plant Mini Kit (Qiagen) according to the manufacturer’s protocols. The RNA quality was assessed with agarose gel electrophoresis. First strand cDNA was synthesized with SuperScriptIII reverse transcriptase (Invitrogen) according to the manufacturer’s instructions using 500 ng of total RNA as a template. The stilbene synthase transcript was amplified with primers STS1_F and STS1_R for first UV-C and CHX experiments (Fig 1). Primers STS2_F and STS2_R were used in all further RT-PCR experiments. Control reactions amplifying the housekeeping gene actin were done with primers Actin_F and Actin_R. PCR reactions were done using Taq DNA polymerase (Roche Life Science or Thermo scientific) according to the manufacturer’s recommendations and using 1 μl of cDNA as a template with the following program: 94°C for 2 min, 24 cycles of 94°C for 30 s, 57°C for 1 min and 72°C for 1 min followed by 72°C for 7 min.

### Luciferase assay

Luciferase activity was measured from Arabidopsis by sampling 1-2 leaves before and after the treatment. Proteins were extracted on ice with 100 μl of modified lux buffer (Nilsson et al., 1992) and microcentrifuge tube pestles. Cell debris was removed by centrifugation for 10 min at +4 °C. Luciferase activity was measured combining by combining 20 μl of sample with 50 μl of enzyme substrate (Luciferase 1000 Assay System, #E4550, Promega), vortexing sample briefly and counting photons for 5 s in the luminometer (Luminoskan TL plus, generation II, Thermo Labsystems, Finland) at room temperature. Total protein concentration was measured from samples with Bradford assay at 595 nm using BSA as standard. Measured luciferase activity was normalized using total protein concentration.

### Thin layer chromatography (TLC)

Needles from 6-week-old seedlings were treated for 15 min of UV-C, at the distance of 20 cm under an uncovered mercury lamp (sterilAir UVC G9) and ground to powder under liquid nitrogen. Metabolites were extracted with 5ml of methanol per gram wet weight, sonicated for 15 min in water bath sonicator (Finnsonic W181) and left at +4°C overnight. Cell debris was removed with centrifugation and 2+2 μl of extracts and a commercial pinosylvin standard (ArboNova) were applied on TLC plates Silica gel 60 F254 (Merck), which were developed with chloroform: ethyl acetate: formic acid (5:4:1) as the mobile phase. Plates were photographed under UV light using 304 nm filter.

### RNA sequencing analysis

RNA-seq libraries were prepared, sequencing done, and data analysis performed as described in Lim et al. (2016, 2021). In short, all transcriptome libraries were mapped against Pinus EST collection version 9.0 (The Gene Index Databases, 2014) using SHRiMP alignment tool (David et al., 2011). Pinus EST collection is composed of tentative consensus (TC) sequences that are assemblies from highly similar sequences from different pine species. For instance, TC154538 for stilbene synthase is 98% identical to the GenBank Scots pine STS mRNA sequence S50350 (Lim et al., 2016). Differential expression analysis was done using edgeR (Robinson et al., 2010) and Bayesian model-based hierarchical clustering using *SplineCluster* (Heard et al., 2006). The RNA-seq reads have been submitted to NCBI under Bioproject PRJNA635551.

### *In silico* promoter analysis, GO enrichment and intersect analysis

The Scots pine stilbene synthase promoter sequence (PST-1) (Preisig-Müller et al.,1999) was submitted to plant promoter analysis database PlantPan 2.0 (Chow et al., 2016) to screen for putative TF binding elements.

All differentially expressed genes from *SplineCluster* analysis in CHX and UV-C treatments were grouped in four patterns according to their expression profiles through time after treatment. GO enrichment analysis was performed for each pattern using the whole annotation of detected ESTs in each treatment as a reference set and the GO categories molecular function, biological process and cellular component were analyzed. GO enrichment analysis was performed using Biological Networks Gene Ontology tool (BiNGO) with hypergeometric test to assess statistical significance of GO terms (Maere et al., 2005). Benjamin and Hochberg false discovery rate (FDR) correction was selected for multiple comparisons with a *p-value* threshold of 0.05.

All differentially expressed transcripts associated with GO terms GO:0003700 (sequence-specific DNA binding transcription factor activity) or GO:0003677 (DNA binding) were extracted from UV-C and CHX RNA-seq data and from previously published TZ/SW (Lim et al. 2016) and wounding (Lim et al., 2021) RNA-seq data. Intersection analysis between differentially expressed genes in CHX, UV-C, TZ/SW and wounding treatments was performed using *venn* R package.

## Acknowledgements

We thank Dr. Matthew Robson and Dr. Titta Kotilainen for measuring the spectral photon irradiance from the UV-C light source used in the experiments. This manuscript has been co-authored by UT-Battelle, LLC, under contract DE-AC05-00OR22725 with the U.S. Department of Energy (DOE). The U.S. government retains and the publisher, by accepting the article for publication, acknowledges that the U.S. government retains a nonexclusive, paid-up, irrevocable, worldwide license to publish or reproduce the published form of this manuscript, or allow others to do so, for U.S. government purposes. DOE will provide public access to these results of federally sponsored research in accordance with the DOE Public Access Plan (http://energy.gov/downloads/doe-public-access-plan).

## Author contributions

T.P. wrote the manuscript; T.P.and K.-J.L designed and carried out the experiments; K.-J.L. M.P. and T.P performed data analysis; L.P. and P.A. were responsible for designing the library phase and conducting SOLiD sequencing; T.H.T., K.K., A.H., and M.V. designed the research; T.H.T. contributed in interpreting the data and revised the manuscript, which all authors read, commented on, and approved.

## References

Bahrami S, Drabløs F (2016) Gene regulation in the immediate-early response process. Advances in Biological Regulation 62: 37–49

Belhadj A, Telef N, Saigne C, Cluzet S, Barrieu F, Hamdi S, Mérillon J (2008) Effect of methyl jasmonate in combination with carbohydrates on gene expression of PR proteins, stilbene and anthocyanin accumulation in grapevine cell cultures. Plant Physiology and Biochemistry 46: 493–499

Belt T, Venäläinen M, Harju A (2021) Non-destructive measurement of Scots pine heartwood stilbene content and decay resistance by means of UV-excited fluorescence spectroscopy. Industrial Crops and Products 164: 113395

Bomal C, Bedon F, Caron S, Mansfield SD, Levasseur C, Cooke JEK, Blais S, Tremblay L, Morency M, Pavy N, Grima-Pettenati J, Séguin A, MacKay J (2008) Involvement of Pinus taeda MYB1 and MYB8 in phenylpropanoid metabolism and secondary cell wall biogenesis: a comparative in planta analysis. J Exp Bot 59: 3925–3939

Bomal C, Duval I, Giguère I, Fortin É, Caron S, Stewart D, Boyle B, Séguin A, MacKay JJ (2013) Opposite action of R2R3-MYBs from different subgroups on key genes of the shikimate and monolignol pathways in spruce. J Exp Bot 65: 495–508

David M, Dzamba M, Lister D, Ilie L & Brudno M (2011) SHRiMP2: sensitive yet practical short read mapping. Bioinformatics 27, 1011–1012

Chiron H, Drouet A, Claudot A, Eckerskorn C, Trost M, Heller W, Ernst D, Sandermann H (2000) Molecular cloning and functional expression of a stress-induced multifunctional O-methyltransferase with pinosylvin methyltransferase activity from Scots pine (Pinus sylvestris L.). Plant Mol Biol 44: 733–745

Chong J, Poutaraud A, Hugueney P (2009) Metabolism and roles of stilbenes in plants. Plant Science 177: 143–155

Chow CN, Zheng HQ, Wu NY, Chien CH, Huang HD, Lee TY, Chiang-Hsieh YF, Hou PF, Yang TY, Chang WC (2016) PlantPAN 2.0: an update of plant promoter analysis navigator for reconstructing transcriptional regulatory networks in plants. Nucleic Acids Res. 4:1154–60.

Chung HS, Koo AJK, Gao X, Jayanty S, Thines B, Jones AD, Howe GA (2008) Regulation and function of Arabidopsis JASMONATE ZIM-Domain genes in response to wounding and herbivory. Plant Physiol 146: 952–964

Clough SJ, Bent AF (1998). Floral dip: a simplified method for Agrobacterium-mediated transformation of Arabidopsis thaliana. Plant J. 16:735–43

Costa MA, Bedgar DL, Moinuddin SGA, Kim K-W, Cardenas CL, Cochrane FC, Shockey JM, Helms GL, Amakura Y, Takahashi H, Milhollan JK, Davin LB, Browse J, Lewis NG (2005). Characterization in vitro and in vivo of the putative multigene 4-coumarate:CoA ligase network in Arabidopsis: syringyl lignin and sinapate/sinapyl alcohol derivative formation. Phytochemistry 66: 2072–2091

Danon A, Rotari VI, Gordon A, Mailhac N, Gallois P (2004) Ultraviolet-C overexposure induces programmed cell death in Arabidopsis, which is mediated by Caspase-like activities and which can be suppressed by caspase inhibitors, p35 and defender against apoptotic death. Journal of Biological Chemistry 279: 779–787

Ehlting J, Büttner D, Wang Q, Douglas CJ, Somssich IE, Kombrink E (1999) Three 4-coumarate:coenzyme A ligases in Arabidopsis thaliana represent two evolutionarily divergent classes in angiosperms. The Plant Journal 19: 9–20

Falcone Ferreyra ML, Rius SP, Casati P (2012) Flavonoids: biosynthesis, biological functions, and biotechnological applications. Front. Plant Sci. 3:222.

Gaid MM, Sircar D, Müller A, Beuerle T, Liu B, Ernst L, Hänsch R, Beerhues L (2012) Cinnamate:CoA ligase initiates the biosynthesis of a benzoate-derived xanthone phytoalexin in Hypericum calycinum cell cultures. Plant Physiology 160: 1267–1280

Gallego F, Fleck O, Li A, Wyrzykowska J, Tinland B (2000) AtRAD1, a plant homologue of human and yeast nucleotide excision repair endonucleases, is involved in dark repair of UV damages and recombination. The Plant Journal 21: 507–518.

Gao C, Xing D, Li L, Zhang L (2008) Implication of reactive oxygen species and mitochondrial dysfunction in the early stages of plant programmed cell death induced by ultraviolet-C overexposure. Planta 227: 755–767

Gentile M, Latonen L, Laiho M (2003) Cell cycle arrest and apoptosis provoked by UV radiation-induced DNA damage are transcriptionally highly divergent responses. Nucleic Acids Research 31:4779–4790

Grimmig B, Schubert R, Fischer R, Hain R, Schreier PH, Betz C, Langebartels C, Ernst D, Sandermann H (1997) Ozone- and ethylene-induced regulation of a grapevine resveratrol synthase promoter in transgenic tobacco. Acta Physiologiae Plantarum 19: 467–474

Harju A, Venäläinen M (2006) Measuring the decay resistance of Scots pine heartwood indirectly by the Folin-Ciocalteu assay. Canadian Journal of Forest Research 36: 1797–1804

Harju A, Venäläinen M, Laakso T, Saranpää P (2009) Wounding response in xylem of Scots pine seedlings shows wide genetic variation and connection with the constitutive defence of heartwood. Tree Physiology 29:19–25

Heard NA, Holmes CC, Stephens DA (2006). A Quantitative study of gene regulation involved in the immune response of Anopheline mosquitoes. Journal of the American Statistical Association 101: 18–29

Höll J, Vannozzi A, Czemmel S, D’Onofrio C, Walker AR, Rausch T, Lucchin M, Boss PK, Dry IB, Bogs J (2013) The R2R3-MYB transcription factors MYB14 and MYB15 regulate stilbene biosynthesis in Vitis vinifera. The Plant Cell 25: 4135–4149

Hovelstad H, Leirset I, Oyaas K, Fiksdahl A (2006) Screening analyses of pinosylvin stilbenes, resin acids and lignans in Norwegian conifers. Molecules 11: 103–114

Jeandet P, Sbaghi M, Bessis R, Meunier P (1995) The potential relationship of stilbene (resveratrol) synthesis to anthocyanin content in grape berry skins. VITIS - Journal of Grapevine Research 34

Jenkins GI (2009) Signal transduction in responses to UV-B radiation. Annu Rev Plant Biol 60: 407–431

Jiang J, Xi H, Dai Z, Lecourieux F, Yuan L, Liu X, Patra B, Wei Y, Li S, Wang L (2019) VvWRKY8 represses stilbene synthase genes through direct interaction with VvMYB14 to control resveratrol biosynthesis in grapevine. J Exp Bot 70:715–729

Klempien A, Kaminaga Y, Qualley A, Nagegowda DA, Widhalm JR, Orlova I, Shasany AK, Taguchi G, Kish CM, Cooper BR, D’Auria JC, Rhodes D, Pichersky E, Dudareva N (2012) Contribution of CoA ligases to benzenoid biosynthesis in Petunia flowers. The Plant Cell 24: 2015–2030

Kodan A, Kuroda H, Sakai F (2002) A stilbene synthase from Japanese red pine (Pinus densiflora): Implications for phytoalexin accumulation and down-regulation of flavonoid biosynthesis. Proceedings of the National Academy of Sciences 99: 3335–3339

Lim K-J, Paasela T, Harju A, Venäläinen M, Paulin L, Auvinen P, Kärkkäinen K, Teeri TH (2016) Developmental changes in Scots pine transcriptome during heartwood formation. Plant Physiology 172: 1403–1417

Lim K-J, Paasela T, Harju A, Venäläinen M, Paulin L, Auvinen P, Kärkkäinen K, Teeri TH (2021) A transcriptomic view to wounding response in young Scots pine stems. Sci Rep 11: 3778

Lorenzo O, Piqueras R, Sánchez-Serrano JJ, Solano R (2003) ETHYLENE RESPONSE FACTOR1 integrates signals from ethylene and jasmonate pathways in plant defense. Plant Cell 15: 165

Mankin SL, Allen GC, Thompson WF (1997) Introduction of a plant intron into the luciferase gene of Photinus pyralis. Plant Mol Biol Rep 15:186–196

Michiels A, Van den Ende W, Tucker M, Van Riet L, Van Laere A (2003) Extraction of high-quality genomic DNA from latex-containing plants. Anal Biochem. 315:85–9

Moore JW, Loake GJ, Spoel SH (2011) Transcription dynamics in plant immunity. The Plant Cell 23: 2809–2820

Nawkar GM, Maibam P, Park JH, Sahi VP, Lee SY, Kang CH (2013) UV-induced cell death in plants. International Journal of Molecular Sciences 14: 1608–1628

Nilsson M, Wikman S, Eklund L (2002) Induction of discolored wood in Scots pine (Pinus sylvestris). Tree Physiol 22: 331–338

Nilsson O, Aldén T, Sitbon F, Little CHA, Chalupa V, Sandber G, Olsson O (1992) Spatial pattern of cauliflower mosaic virus 35S promoter-luciferase expression in transgenic hybrid aspen trees monitored by enzymatic assay and non-destructive imaging. Transgenic Research 1:209–220

O’Donnell PJ, Calvert C, Atzorn R, Wasternack C, Leyser HMO, Bowles DJ (1996) Ethylene as a signal mediating the wound response of tomato plants. Science 274: 1914

Paasela T, Lim K, Pietiäinen M, Teeri TH (2017) The O-methyltransferase PMT2 mediates methylation of pinosylvin in Scots pine. New Phytol 214: 1537–1550

Patzlaff A, McInnis S, Courtenay A, Surman C, Newman LJ, Smith C, Bevan MW, Mansfield S, Whetten RW, Sederoff RR, Campbell MM (2003) Characterisation of a pine MYB that regulates lignification. The Plant Journal 36: 743–754

Penninckx IAMA, Thomma BPHJ, Buchala A, Métraux J, Broekaert WF (1998) Concomitant activation of jasmonate and ethylene response pathways is required for induction of a plant Defensin gene in Arabidopsis. The Plant Cell 10: 2103–2113

Preisig-Müller R, Schwekendiek A, Brehm I, Reif H, Kindl H (1999) Characterization of a pine multigene family containing elicitor-responsive stilbene synthase genes. Plant Mol Biol 39: 221–229

Rodríguez-Concepción M, Boronat A (2002) Elucidation of the methylerythritol phosphate pathway for isoprenoid biosynthesis in bacteria and plastids. A metabolic milestone achieved through genomics. Plant Physiology 130: 1079–1089

Robinson MD, McCarthy DJ & Smyth GK (2010) edgeR: a bioconductor package for differential expression analysis of digital gene expression data. Bioinformatics 26, 139–140

Schanz S, Schröder G, Schröder J (1992) Stilbene synthase from Scots pine (Pinus sylvestris). FEBS Lett 313: 71–74

Schoeppner A, Kindl H (1979) Stilbene synthase (pinosylvine synthase) and its induction by ultraviolet light. FEBS Lett 108: 349–352

Schubert R, Fischer R, Hain R, Schreier PH, Bahnweg G, Ernst D, Sandermann Jr H (1997) An ozone-responsive region of the grapevine resveratrol synthase promoter differs from the basal pathogen-responsive sequence. Plant Mol Biol 34: 417–426

Schwekendiek A, Pfeffer G, Kindl H (1992) Pine stilbene synthase cDNA, a tool for probing environmental stress. FEBS Lett 301: 41–44

Suzuki M, Nakabayashi R, Ogata Y, Sakurai N, Tokimatsu T, Goto S, Suzuki M, Jasinski M, Martinoia E, Otagaki S, Matsumoto S, Saito K, Shiratake K (2015) Multiomics in grape berry skin revealed specific induction of the stilbene synthetic pathway by ultraviolet-C irradiation. Plant Physiology 168: 47–59

Tassoni A, Fornalè S, Franceschetti M, Musiani F, Michael AJ, Perry B, Bagni N (2005) Jasmonates and Na-orthovanadate promote resveratrol production in Vitis vinifera cv. Barbera cell cultures. New Phytol 166: 895–905

Tenhaken R, Rübel C (1998) Induction of alkalinization and an oxidative burst by low doses of cycloheximide in soybean cells. Planta 206: 666–672

Tullai JW, Schaffer ME, Mullenbrock S, Sholder G, Kasif S, Cooper GM (2007) Immediate-early and delayed primary response genes are distinct in function and genomic architecture. Journal of Biological Chemistry 282: 23981–23995

Usami S, Banno H, Ito Y, Nishihama R, Machida Y (1995) Cutting activates a 46-kilodalton protein kinase in plants. Proc Natl Acad Sci U S A 92: 8660–8664

Wang D, Jiang C, Liu W, Wang Y (2019) The WRKY53 transcription factor enhances stilbene synthesis and disease resistance by interacting with MYB14 and MYB15 in Chinese wild grape, J. Exp. Bot. 71: 3211–3226

Vannozzi A, Dry IB, Fasoli M, Zenoni S, Lucchin M (2012) Genome-wide analysis of the grapevine stilbene synthase multigenic family: genomic organization and expression profiles upon biotic and abiotic stresses. BMC Plant Biology 12: 130

Vannozzi A, Wong DCJ, Höll J, Hmmam I, Matus JT, Bogs J, Ziegler T, Dry I, Barcaccia G, Lucchin M (2018). Combinatorial regulation of stilbene synthase genes by WRKY and MYB transcription factors in grapevine (Vitis vinifera L.), Plant and Cell Physiology, 59: 1043–1059

Venäläinen M, Harju AM, Saranpää P, Kainulainen P, Tiitta M, Velling P (2004) The concentration of phenolics in brown-rot decay resistant and susceptible Scots pine heartwood. Wood Sci Technol 38: 109–118

Wong DCJ, Schlechter R, Vannozzi A, Höll J, Hmmam I, Bogs J, Tornielli GB, Castellarin SD, Matus JT (2016) A systems-oriented analysis of the grapevine R2R3-MYB transcription factor family uncovers new insights into the regulation of stilbene accumulation. DNA Research 23: 451–466

Xi H, Ma L, Liu G, Wang N, Wang J, Wang L, Dai Z, Li S, Wang L (2014) Transcriptomic analysis of grape (Vitis vinifera L.) leaves after exposure to Ultraviolet C Irradiation. PLOS ONE 9: e113772

Xu Y, Chang P, Liu D, Narasimhan ML, Raghothama KG, Hasegawa PM, Bressan RA (1994) Plant defense genes are synergistically induced by ethylene and methyl jasmonate. Plant Cell 6: 1077

